# Allele-specific gene editing rescues pathology in a human model of Charcot-Marie-Tooth disease type 2E

**DOI:** 10.1101/2021.06.09.447796

**Authors:** Carissa M. Feliciano, Kenneth Wu, Hannah L. Watry, Chiara B.E. Marley, Gokul N. Ramadoss, Hana Y. Ghanim, Angela Z. Liu, Lyandysha V. Zholudeva, Todd C. McDevitt, Mario A. Saporta, Bruce R. Conklin, Luke M. Judge

## Abstract

Many neuromuscular disorders are caused by dominant missense mutations that lead to dominant-negative or gain-of-function pathology. This category of disease is challenging to address via drug treatment or gene augmentation therapy because these strategies may not eliminate the effects of the mutant protein or RNA. Thus, effective treatments are severely lacking for these dominant diseases, which often cause severe disability or death. The targeted inactivation of dominant disease alleles by gene editing is a promising approach with the potential to completely remove the cause of pathology with a single treatment. Here, we demonstrate that allele-specific CRISPR gene editing in a human model of axonal Charcot-Marie-Tooth (CMT) disease rescues pathology caused by a dominant missense mutation in the neurofilament light chain gene (*NEFL*, CMT type 2E). We utilized a rapid and efficient method for generating spinal motor neurons from human induced pluripotent stem cells (iPSCs) derived from a patient with CMT2E. Diseased motor neurons recapitulated known pathologic phenotypes at early time points of differentiation, including aberrant accumulation of neurofilament light chain protein in neuronal cell bodies. We selectively inactivated the disease *NEFL* allele in patient iPSCs using Cas9 enzymes to introduce a frameshift at the pathogenic N98S mutation. Motor neurons carrying this allele-specific frameshift demonstrated an amelioration of the disease phenotype comparable to that seen in an isogenic control with precise correction of the mutation. Our results validate allele-specific gene editing as a therapeutic approach for CMT2E and as a promising strategy to silence dominant mutations in any gene for which heterozygous loss-of-function is well tolerated. This highlights the potential for gene editing as a therapy for currently untreatable dominant neurologic diseases.

## INTRODUCTION

Charcot-Marie-Tooth disease (CMT), also known as hereditary motor and sensory neuropathy, is one of the most common inherited neurological disorders, affecting approximately 1 in 2,500 people in the United States (Reilly et al., 2011). Patients present with muscle weakness and variable sensory symptoms in the distal limbs, which typically progress in a length-dependent manner, affecting the most distal limbs first. CMT can be further classified by the predominantly affected cell type, by clinical electrophysiological testing, and by underlying genetic cause (Rossor et al., 2015). The demyelinating forms of the disease (CMT1, autosomal dominant and CMT4, autosomal recessive) primarily affect Schwann cells and are characterized by a decrease in nerve conduction velocity (Brennan et al., 2015). The axonal forms of the disease (CMT2, dominant and recessive) are characterized by pathology intrinsic to motor neurons (Luigetti et al., 2016). The genetic causes of CMT are remarkably diverse, with causative mutations in over 80 genes, and a diverse set of individual mutations reported for many of those genes (Timmerman et al., 2014; DiVincenzo et al., 2014; Pareyson et al., 2017).

Axonal CMT is predominantly caused by autosomal dominant missense mutations, which occur in more than 20 genes with diverse functions (Rossor et al., 2015; Pareyson et al., 2017). A notable example is CMT2E, caused by mutations in the *NEFL* gene, which encodes the neurofilament light chain (NF-L) protein. Neurofilaments are neuron-specific intermediate filaments composed of multiple subunits, including NF-L, that contribute to the radial growth of axons and the maintenance of axonal diameter and function (Szaro and Strong, 2010; Yuan et al., 2017). A multitude of missense mutations throughout the *NEFL* coding sequence cause autosomal dominant CMT2E (Stone et al., 2019). By contrast, loss-of-function mutations in *NEFL* are recessive, and only cause CMT when homozygous—the heterozygous carriers are neurologically normal (Abe et al., 2009; Yum et al., 2009; Fu and Yuan, 2018). These loss-of-function mutations demonstrate that a single functional *NEFL* allele is adequate for normal neurological function and suggest that dominant missense mutations in *NEFL* act via dominant-negative or pathologic gain-of-function mechanisms. The N98S mutation is a prime example of the latter, as a single mutant allele leads to the aberrant accumulation of neurofilaments in neuronal cell bodies and causes a severe and early-onset motor neuropathy in humans and mice (Adebola et al., 2015; Saporta et al., 2015; Zhao et al., 2017; Jordanova et al., 2003; Lancaster et al., 2018).

There is currently no effective treatment for any form of CMT. Viral gene replacement therapy for the motor neuron disease spinal muscular atrophy (SMA) has shown great promise and could represent a viable strategy for the rare recessive forms of CMT2 (Mendell et al., 2017). However, dominant CMT2 likely requires an alternate approach as simply increasing expression of a wild-type gene may not overcome the pathologic effects of the mutant allele, which will continue to be expressed. Advances in gene-editing technologies create an opportunity to develop therapies that interrupt the pathologic processes causing dominant CMT2, and other dominant neurodegenerative disorders, at its genetic source (Porteus, 2019). Since people with a single functional *NEFL* allele have normal neurological function, we reasoned that selective inactivation of dominant missense mutations in *NEFL* via CRISPR-Cas9 gene editing would be therapeutic. We tested this hypothesis using an established human induced pluripotent stem cell (iPSC) model of CMT2E with the N98S mutation in *NEFL* (Saporta et al., 2015).

Here, we report a gene-editing strategy specific for the N98S mutation that efficiently and selectively reduced expression of the mutant allele in CMT2E motor neurons. Edited neurons displayed phenotypes that were similar to unrelated and isogenic controls, indicating that the editing strategy effectively rescued disease-associated pathology. This is the first reported application of gene editing for axonal CMT and provides proof-of-principle for strategies that could be applied to other dominant-negative inherited neurodegenerative diseases.

## MATERIALS AND METHODS

### iPSC culture

The healthy control iPSC line (WTC) used in these studies is an extensively characterized line from a healthy individual (Kreitzer et al., 2013) that is the parental line for the Allen Institute Cell Collection (http://www.allencell.org/). The CMT2E iPSC line was derived from an individual with the N98S mutation in *NEFL* causing childhood onset CMT and has been previously characterized (Saporta et al., 2015; Maciel et al., 2020). iPSCs were cultured in Stemfit Basic02 (Ajinomoto) on plates coated with matrigel (Corning, 356231) at 37 °C, 5% CO_2_, 85% humidity. iPSC cultures were passaged every 3-4 days with Accutase (Stem Cell Technologies, 07920) or ReLeSR (Stem Cell Technologies, 05872) into Stemfit supplemented with 10 μm Y-27632 (SelleckChem, S1049).

### Generating transgenic iPSC lines with inducible transcription factors

The iPSCs were engineered to contain a 13kb vector containing the human transcription factors *NGN2*, *ISL1*, and *LHX3* (hNIL) under the control of doxycycline (Fernandopulle et al., 2018). To knock-in the hNIL construct into safe harbor loci, 1 x 10^6^ cells were added to 100 μL P3 buffer with 6 μg of the hNIL donor plasmid, and 3 μg of each paired TALEN targeting either AAVS1 or CLYBL (Addgene, 59025/59026 or 62196/62197, respectively). Cells were transfected using the P3 Primary Cell 4D-Nucleofector X Kit L (Lonza, V4XP-3024) with pulse code DS138. After nucleofection, cells incubated for 5 minutes at room temperature and were then plated in Stemfit media with 10 μm Y-27632 in a serial dilution into 5 wells of a 6-well plate. Non-transfected cells were plated in the sixth well. After 2-3 days post-nucleofection, cells were fed with Stemfit with Puromycin or G418 (depending on which donor vector was used) and 10 μM Y-27632. Cells were kept on antibiotic selection until all cells in the control wells were dead and only red fluorescent colonies remained in the experimental wells. Fluorescent, antibiotic resistant colonies were then manually picked and transferred to individual wells of a 48-well plate.

DNA from each clone was extracted using the DNeasy Blood and Tissue Kit (Qiagen, 69506). PCRs were performed with primers spanning the 5’ and 3’ junctions of the integration, with one primer annealing within the construct and the other outside of the corresponding homology arm. A third PCR amplifying the intact safe harbor locus was performed to determine whether the clone harbored a heterozygous or homozygous insertion. PCR primer sequences are listed in Supplementary Table 1. For the CMT2E line, we inserted hNIL in the AAVS1 locus (Supplementary Fig. S1) and obtained a previously published WTC line with hNIL inserted in the AAVS1 locus as a control (Shi et al., 2018). We also independently inserted hNIL in the CLYBL locus of the same parental WTC line (Supplementary Fig. S1). When performing neuronal differentiation, we found equivalent results with the AAVS1 and CLYBL versions of the WT line. For comparison with the CMT2E-hNIL-AAVS1 line, we used WTC-hNIL-AAVS1 for all experiments, with the exception of manual image analysis shown in Figure 2, in which case we used the WTC-hNIL-CLYBL line.

### CRISPR/Cas9 gene-editing analysis

To prepare the Sp.HiFi Cas9 RNP complex, 240 pmol of guide RNA (IDT) was mixed with 120 pmol of Hi-Fi SpCas9 protein (Macrolab) and incubated for 30 min at room temperature. For the Sa.KKH Cas9 RNP complex, 240 pmol of guide RNA (Synthego) was mixed with 120 pmol of Sa.KKH Cas9 protein (Macrolab). The gRNA sequences are listed in Supplementary Table 2. After dissociation with Accutase, 3.5 x 10^5^ cells were resuspended in P3 buffer and mixed with the RNP complex. The iPSCs were transfected using the P3 Primary Cell 4D-Nucleofector X Kit S (Lonza, V4XP-3032) with pulse code DS138. After nucleofection, cells were incubated for 5 min at room temperature and then plated in Stemfit with 10 μM Y-27632 (SelleckChem). Genomic DNA was extracted from edited and unedited cells 3 days post nucleofection using the DNeasy Blood and Tissue Kit (Qiagen). PCRs were performed with primers spanning the gRNA binding site (Supplementary Table 1), and the PCR products were sequenced. The indel frequency at the cut site was determined using Synthego ICE software, which compares the Sanger sequencing traces from the edited and unedited populations (Supplementary Fig. S2).

### Isolation of clonal edited lines

To generate the N98S-fs line, 3.5 x 10^5^ cells were transfected with RNP as described above. After nucleofection, 0.1 x 10^5^ cells were seeded onto a 10 cm plate in Stemfit with 10 μM Y-27632. After 5 days post-nucleofection, colonies were manually picked and transferred to individual wells of a 48-well plate. DNA was extracted from the cells using QuickExtract DNA Extract Solution (Lucigen, QE9050). Clones were genotyped by PCR and Sanger Sequencing (Supplementary Fig. 3B).

To generate the N98S-corrected line, 3.5 x 10^5^ cells were nucleofected with RNP and 50 pmol of donor DNA (IDT, Supplementary Table 1). To enrich for the HDR event and isolate a clone with the desired edit, allele-specific ddPCR and sib-selection were performed ((Miyaoka et al., 2014), Supplementary Fig. 3C). After nucleofection, cells were plated at 2.5 x 10^3^ cells per well in a 96-well plate. DNA was extracted from the cells using QuickExtract and analyzed by allele-specific ddPCR to measure HDR efficiency. The well with the highest HDR efficiency was replated at 10 cells per well in a 96-well plate for an additional round of enrichment. From this round, the well with the highest HDR efficiency (~11.4%) was replated sparsely into a 10 cm plate for manual clone picking. The clones were genotyped by allele-specific ddPCR and further validated by PCR and Sanger Sequencing (Supplementary Fig. 3D). The clonal cell line with the desired HDR event underwent an additional round of manual clone picking.

### i^3^LMN differentiation

iPSCs were differentiated into i^3^LMNs as described in (Fernandopulle et al., 2018), with the following modifications. On day 3, cells used for immunocytochemistry were seeded at 5-8 x 10^4^ cells per cm^2^ onto poly-D-lysine (PDL) (Sigma, P7405) and laminin-coated 12 mm glass coverslips in 24-well plates or PDL and laminin-coated clear bottom imaging 96-well plates. Cells used for RNA and protein assays were seeded at 1-1.5 x 10^5^ cells per cm^2^ onto PDL and laminin-coated 6-well plates. On day 4, the media was removed and replaced with fresh Neural Induction Media (NIM) supplemented with B-27 (Gibco, 17504-044), CultureOne (Gibco, A33202-01), 1 μg/mL laminin, 20 ng/mL BDNF (PeproTech, 450-02), 20 ng/mL GDNF (PeproTech, 450-10), and 10 ng/mL NT3 (PeproTech, 450-03). On day 7, a half volume of media was aspirated and replaced with fresh NIM supplemented with B-27, CultureOne, 1 μg/mL laminin, 40 ng/mL BDNF, 40 ng/mL GDNF, and 20 ng/mL NT3. On day 10, a half volume of media was aspirated and replaced with fresh Neural Maintenance Media (NMM) supplemented with B-27, CultureOne, 1 μg/mL laminin, 40 ng/mL BDNF, 40 ng/mL GDNF, and 20 ng/mL NT3.

### Immunofluorescent staining

i^3^LMNs used for immunofluorescent staining were seeded onto glass coverslips or clear bottom imaging 96-well plates on day 3 of the differentiation, which then proceeded as previously described. For fixation, an equivalent volume of 4% paraformaldehyde (PFA, Alfa Aesar, J61899AK) was added directly to the cell culture media. Cells were incubated in PFA at room temperature for 20 min, then washed with PBS with 0.1% Triton-X (Sigma, X100) (PBS-T) once quickly, followed by two 15-min washes. The cells were blocked and permeabilized for one hour at room temperature with 5% BSA (Sigma, A4503) in PBS-T. The cells were incubated in primary antibodies at appropriate dilutions (Supplementary Table 3) in blocking buffer at 4°C overnight. The following day, the cells were washed with PBS-T at room temperature once quickly, followed by three 10-min washes. The cells were incubated in fluorescent conjugated secondary antibodies diluted at 1:500 in blocking buffer for 1 hour at room temperature. The cells were washed with PBS-T once quickly, followed by three 10-min washes. The first 10-min wash contained DAPI dye (Invitrogen, D1306). The coverslips were then mounted onto a slide with VECTASHIELD HardSet Mounting Media (Vector Laboratories, H-1400-10). Images were taken on a Keyence BZ-9000 Fluorescence Microscope.

### RNA expression assays

i^3^LMNs used for RNA-based assays were collected on day 7 of the differentiation. RNA was extracted from iPSCs and i^3^LMNs using the Quick-RNA Miniprep Kit (Zymo Research, R1055), according to the manufacturer’s instructions. RNA concentrations were quantified using the NanoDrop spectrophotometer. Reverse transcription was performed using the iScript cDNA Synthesis Kit (Bio-Rad, 1708891).

To determine mRNA expression for *HB9*, *CHAT*, and *NEFL*, we used the respective TaqMan gene expression assays (ThermoFisher) containing a FAM-labeled probe. All reactions included a GAPDH gene expression assay (Bio-Rad) containing a HEX-labeled probe, which was used as an internal control to normalize levels of mRNA. 5 ng cDNA were used in the 25 μL reactions measuring *HB9* and *CHAT* expression. 0.25 ng cDNA were used in the 25 μL reactions measuring *NEFL* expression. The gene expression assay IDs are listed in Supplementary Table 4. To determine allele-specific *NEFL* mRNA expression, we used a TaqMan SNP genotyping assay targeting the rs79736124 SNP in the 3’ UTR of *NEFL* (ThermoFisher, C_105316276_10). 0.25 ng cDNA were used in each 25 μL ddPCR reaction.

Each 25 μL ddPCR reaction contained 12.5 μL 2x Supermix for Probes (no dUTP) (Bio-Rad Laboratories, #186-3024), cDNA, primer and probe mix(es), and water to 25 μL. Droplets were generated using 20 μL of reaction mixture and 70 μL of oil with the QX200 Droplet Generator. Droplets were transferred to a 96-well PCR plate, sealed, and run on a C1000 Thermal Cycler with a deep well block (Bio-Rad Laboratories). All ddPCR reactions were run under the following thermal cycling conditions: 1) 95 °C for 10 min; 2) 94 °C for 30 s; 3) 58 °C for 1 min; 4) steps 2 and 3 repeat 39 times; 5) 98 °C for 10 min.

All ddPCR runs were analyzed using the Bio-Rad QuantaSoft Pro Software. The ratio of FAM-positive droplets to HEX positive-droplets was used to calculate the gene expression levels for *HB9*, *CHAT*, and *NEFL*. Fractional abundance of FAM and VIC-positive droplets was used to calculate the percentage of wild-type and mutant *NEFL* mRNA, respectively.

### Western blots

i^3^LMNs were harvested on day 10 of the differentiation and lysed in RIPA buffer (ThermoFisher, 89901) with Protease Inhibitor (MedChemExpress, HY-K0010). The samples were incubated for 15 min on ice and sonicated for 8 sec using a probe sonicator. The samples were then incubated for 10 min on ice and centrifuged at 10,000 rpm for 10 min at 4°C. The supernatant of each sample was transferred to a clean tube. The protein was quantified using the Bio-Rad Protein Assay Dye Reagent Concentrate (Bio-Rad, 5000006).

Samples containing 18 μg of total protein were mixed with NuPAGE Sample Reducing Agent (ThermoFisher, NP0009) and NuPAGE LDS Sample Buffer (ThermoFisher, NP0008) and incubated at 70°C for 10 min. The reduced samples were electrophoresed in a 4-12% Bis-Tris gel. Protein was transferred onto an iBlot nitrocellulose membrane (ThermoFisher, IB301002) using the iBlot system (ThermoFisher). The membrane was incubated in Odyssey Blocking Buffer (Li-Cor, 927-4000) and PBS at a 1:1 ratio (blocking buffer) for 1 hour at room temperature. Primary antibodies at appropriate dilutions (Supplementary Table 3) were prepared in blocking buffer with 0.1% Tween 20 (Sigma, 9005-64-5). The membrane was incubated in the primary antibody solution overnight at 4°C. The following day, the membrane was washed with PBS with 0.1% Tween 20 (PBS-T) once quickly and with TBS (Corning, 46-012-CM) with 0.1% Tween 20 (TBS-T) three times for 10 min each. Fluorescent conjugated secondary antibodies at appropriate dilutions were prepared in blocking buffer. The membrane was incubated in the secondary antibody solution for 1 hour at room temperature. The membrane was then washed with TBS-T 3 times for 10 min each and imaged on the Li-Cor Odyssey Fc Imaging System. Protein band intensity was quantified using ImageJ software. NF-L levels were normalized to GAPDH by dividing NF-L signal intensity by GAPDH signal intensity for each biological replicate.

### Image analysis

To assess the accumulation of NF-L in the cell bodies of i^3^LMNs, the i^3^LMNs were fixed and stained with anti-HB9 antibody, anti-NF-L antibody, and DAPI. Ten images were taken per sample on a Keyence BZ-9000 Fluorescence Microscope. The cell bodies of i^3^LMNs were then segmented manually, or automatically using CellProfiler. Before manual segmentation, the images were renamed with alphanumeric codes to allow for blinded analysis. Using ImageJ software, the cell bodies of HB9-positive i^3^LMNs were manually segmented by a blinded observer. Fluorescence intensity was measured in resulting regions of interest from the green channel to measure NF-L intensity within each cell body. To allow for automatic segmentation in later experiments, we designed a CellProfiler pipeline to define cell bodies with HB9-positive nuclei and measure the average NF-L intensity of each cell body (Supplementary Fig. S5). For each sample, the average NF-L intensity in the cell bodies was calculated by averaging the intensity values of all the cell bodies across 10 images.

### NF-L ELISA assay

i^3^LMNs used for the NF-L ELISA Assay were seeded at 30,000 cells per well in a 96-well plate on day 3 of the differentiation. On day 11, a full media change was performed. On day 14, the media from each well was collected and stored at −80°C until the assay was performed. The samples were diluted with water at a 1:25 ratio before quantification. NF-L levels in the media were quantified using the NF-light ELISA kit (Uman Diagnostics, 10-7001).

On day 14, the cells were imaged using the Incucyte S3 Live-Cell Analysis System. Three images per well were analyzed using CellProfiler software to determine the average neurite area (Supplementary Fig. S6). NF-L levels were normalized to the neurite area by dividing the NF-L concentration by the average neurite area for each well.

### Multi-electrode array analysis

iPSCs were differentiated into i^3^LMNs as described in (Fernandopulle et al., 2018), with the following modifications. On day 3, cells were seeded at 20,000 cells per well in 10 μL of Neural Induction Media (NIM) supplemented with 10 μM Y-27632, 2 μg/mL doxycycline, 40 mM BrdU, and 2 mM Compound E onto polyethylenimine (PEI) (Thermofisher, AC178571000) and laminin-coated 24-well CytoView MEA plates (Axion Biosystems, M384-tMEA-24W). After the cells were incubated for 20 minutes at 37°C, 500 μL of complete media was gently added to each well. On day 4, the media was removed and replaced with fresh NIM supplemented with B-27, CultureOne, 1 μg/mL laminin, 20 ng/mL BDNF, 20 ng/mL GDNF, and 10 ng/mL NT3. On day 7, a half volume of media was aspirated and replaced with fresh NIM supplemented with B-27, CultureOne, 1 μg/mL laminin, 40 ng/mL BDNF, 40 ng/mL GDNF, and 20 ng/mL NT3. Starting on day 10, every 3 days, a half volume of media was aspirated and replaced with BrainPhys media (StemCell, 05791) supplemented with B-27, CultureOne, 1 μg/mL laminin, 40 ng/mL BDNF, 40 ng/mL GDNF, and 20 ng/mL NT3. Spontaneous neural activity was measured for 15 minutes using the Axion Biosystem Maestro Edge microelectrode array (MEA) systems. Data was acquired at 12.5 kHz, and spikes were detected in the AxIS Navigator software with an adaptive threshold crossing set to 6 times the standard deviation of the estimated noise for each electrode. The electrical activity was analyzed using weighted mean firing rate to exclude non-active electrodes for the entire 15 minute recording.

## RESULTS

### Rapid development of CMT2E phenotypes in iPSC-derived motor neurons

Multiple groups have demonstrated that the *NEFL* N98S mutation leads to a pathologic accumulation of NF-L in the cell bodies of human and mouse neurons in vitro and in vivo (Adebola et al., 2015; Saporta et al., 2015; Zhao et al., 2017). However, previous methods used to identify phenotypes in motor neurons from CMT2E patient-derived iPSCs were time consuming, often taking ~6 weeks in total (Saporta et al., 2015). To overcome this challenge, we engineered patient-derived iPSCs with integrated, inducible, and isogenic (i^3^) transcription factors in order to rapidly and efficiently differentiate them into lower motor neurons (i^3^LMNs) (Fernandopulle et al., 2018). We inserted a doxycycline-inducible cassette containing *NGN2*, *ISL1*, and *LHX3* into a safe-harbor locus of a previously characterized CMT2E iPSC line with the *NEFL* N98S mutation, referred to here as N98S (Saporta et al., 2015; Maciel et al., 2020). For comparison, we used healthy control (WT) iPSC lines containing the same doxycycline-inducible cassette, as previously described (Shi et al., 2018; Fernandopulle et al., 2018). At day 7 of the differentiation, i^3^LMNs derived from the WT and N98S lines expressed neuronal markers, including neurofilaments, β-III tubulin, and the motor-neuron transcription factor HB9 (*MNX1* gene) (Fig. 1A). Consistent with these results, gene expression analysis by reverse transcriptase-droplet digital polymerase chain reaction (RT-ddPCR) demonstrated that WT and N98S i^3^LMNs expressed *MNX1* and the more mature motor neuron marker choline acetyltransferase (*CHAT*) at day 7 of the differentiation (Fig. 1B). To further verify the functionality of our patient-derived i^3^LMNs, we performed electrophysiological analysis using a multi-electrode array (MEA) device. We detected rare spontaneous action potentials on day 7 that progressively increased in frequency for both cell lines (Fig. 1C).

**Figure 1:**
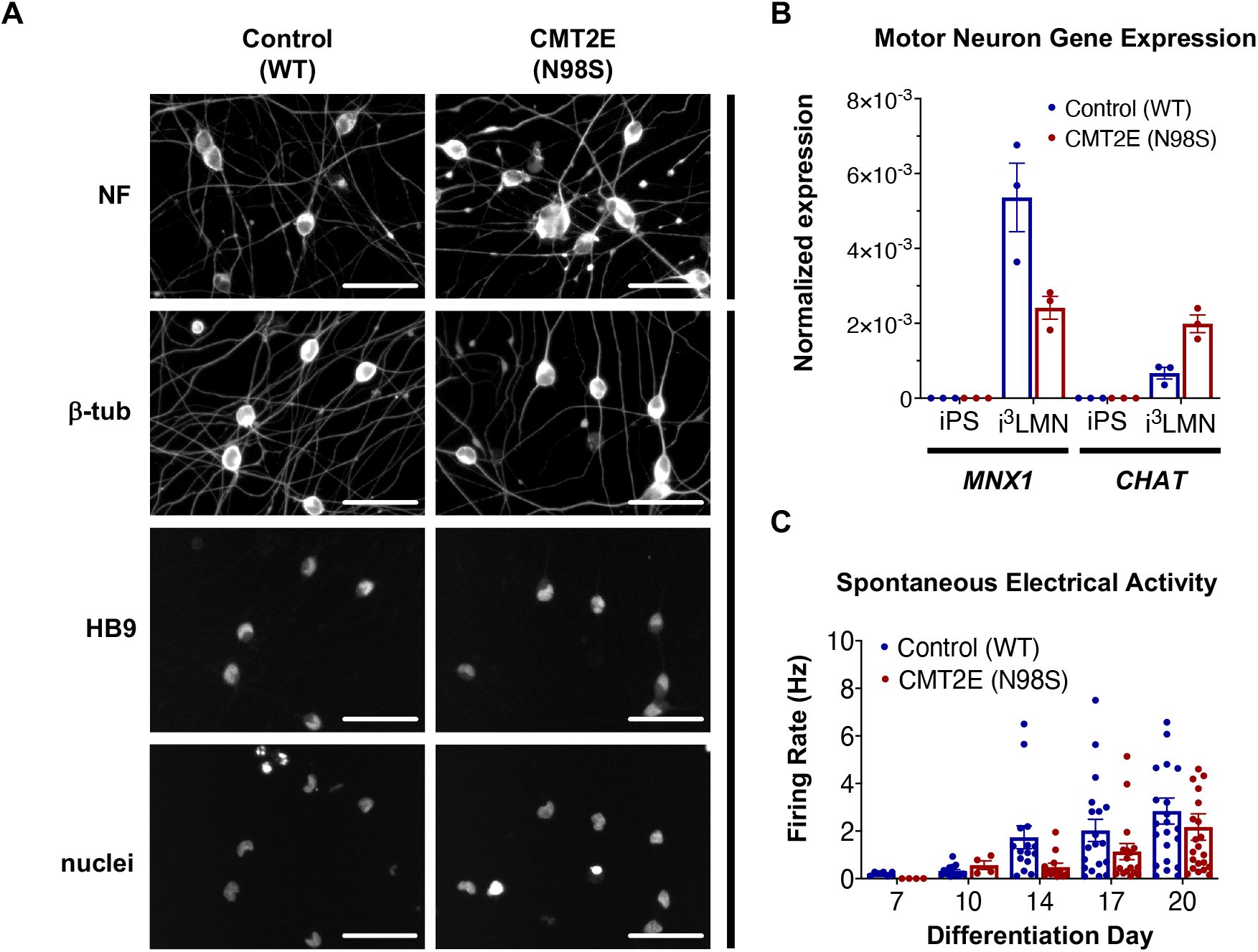
Generation of CMT2E (*NEFL*-N98S) motor neurons. (A) Expression of neuronal markers by immunofluorescent staining in day-7 i^3^LMNs derived from healthy control and CMT2E patient iPSC lines. Lower panels are separate RBG channels from the same images, as indicated by a vertical bar. Scale bars = 50 μM. NF = neurofilaments (SMi^3^3 antibody), β-tub = βIII-tubulin, DAPI stain for nuclei. (B) Total mRNA isolated from undifferentiated iPSCs and day-7 i^3^LMNs was assayed by RT-ddPCR for expression of motor neuron specific genes *MNX1* and *CHAT*, normalized to *GAPDH*. (C) WT and N98S i^3^LMN were seeded on an MEA plate on day 3 of differentiation and spontaneous electrical activity was recorded starting day 7 and every few days until day 20. Graph represents serial measurements of the mean firing rate (frequency of action potentials) from replicate wells. For all graphs individual replicate data points are shown with bars representing mean +/− S.E.M.

To confirm that we could detect disease-relevant phenotypes at early time points in our differentiation, we fixed WT and N98S i^3^LMNs and stained them for NF-L and HB9 using commercially available antibodies (Fig. 2A,B). We observed dense accumulation of NF-L staining in N98S i^3^LMN cell bodies as early as day 7 of differentiation (Fig. 2B). For quantification, NF-L intensity in the cell bodies of HB9-positive cell bodies was measured in a blinded fashion. When examining the distribution of NF-L intensities across all segmented neurons, we observed that a subset of N98S i^3^LMNs exhibited wild-type levels of NF-L in cell bodies but that there was a marked shift toward high-intensity NF-L staining, consistent with a mis accumulation phenotype (Fig. 2C). To further validate this finding, we compared mean NF-L intensities in multiple independent i^3^LMN preparations. We observed a greater than 2-fold increase in mean NF-L intensity in the cell bodies of N98S i^3^LMNs compared to WT i^3^LMNs (Fig. 2D). Thus, we were able to replicate the previously described phenotype, but at a significantly earlier time point compared to other iPSC differentiation methods. We anticipated that this finding would facilitate relatively rapid and sensitive measurement of phenotypic rescue in subsequent gene-editing experiments.

**Figure 2:**
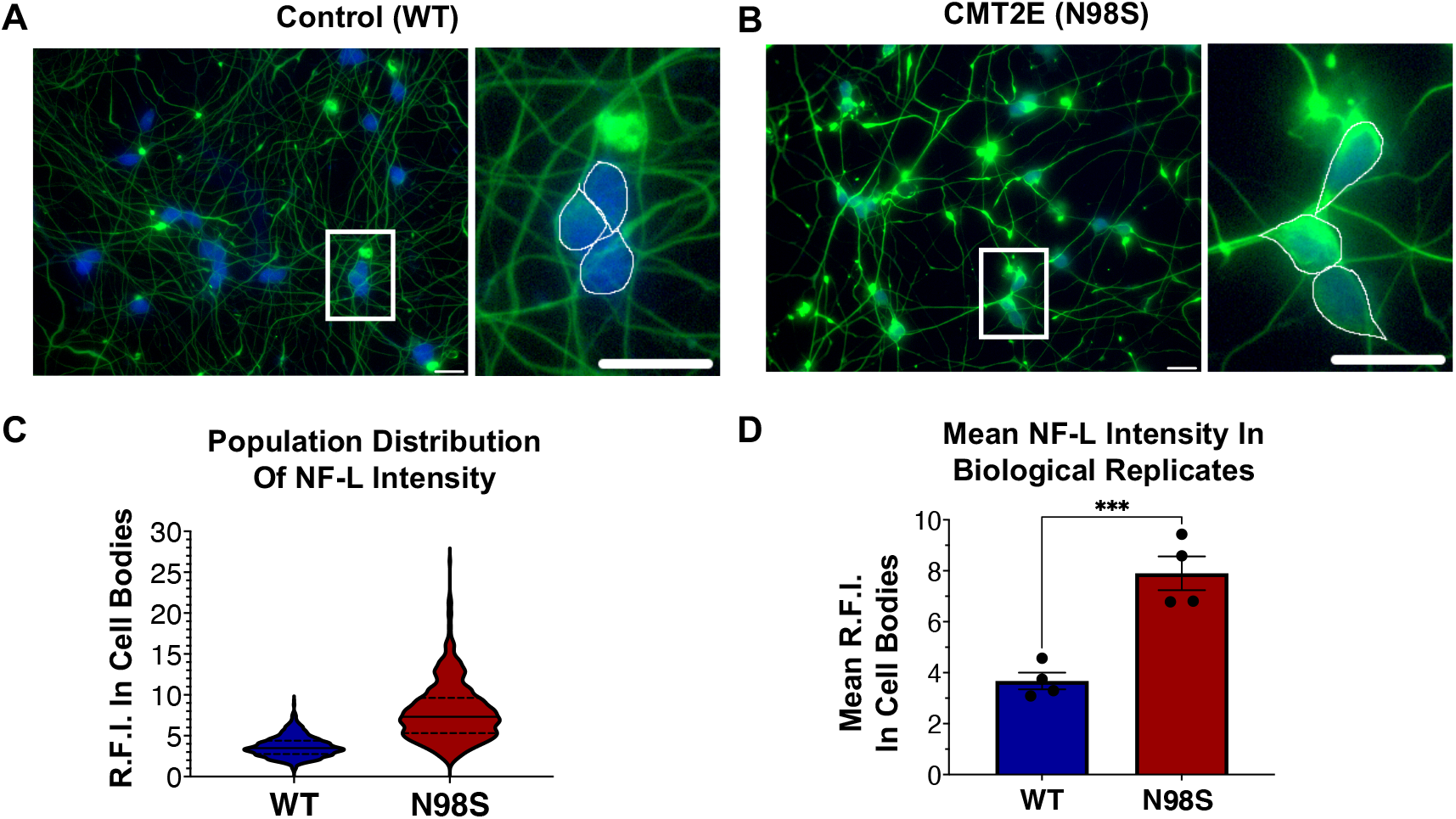
Pathologic accumulation of NF-L protein in the cell body of *NEFL*-N98S motor neurons. (A,B) Representative images of day-7 WT (A) and N98S (B) i^3^LMNs stained with anti-NF-L (green) and anti-HB9 (blue) antibodies. Boxed regions are shown in larger scale to the right of each image with manual cell body segmentation superimposed. Scale bars = 25 μM. (C) Distribution of NF-L fluorescence intensity in individual HB9+ cell bodies after manual segmentation. (D) Quantification of mean NF-L fluorescence intensity from four biological replicates per cell line, demonstrating significantly increased intensity in N98S neurons. Points represent mean of > 160 neurons quantified in each sample population, bars represent mean of all replicates +/− S.E.M., *** p < 0.001 by t-test.

### Allele-specific editing to model therapeutic strategies in CMT2E iPSCs

To enable allele-specific silencing of the N98S allele, we designed guide RNAs targeting the N98S sequence (Fig. 3A) for engineered Cas9 enzyme variants derived from *streptococcus pyogenes* (Sp.HiFi) and *staphylococcal aureus* (Sa.KKH) (Kleinstiver et al., 2015; Vakulskas et al., 2018). We transfected both N98S and WT iPSCs with the resulting Cas9/gRNA ribonucleoprotein (RNP) complexes and performed ICE analysis to measure the generation of insertions and deletions (indels) in the target exon. In N98S iPSCs transfected with Sp.HiFi RNP, approximately 45% of *NEFL* alleles sequenced had indels at the target site (Fig. 3B). This is very close to the 50% maximum expected if all N98S alleles in the heterozygous cell population had been edited. In contrast, in WT iPSCs transfected with Sp.HiFi RNP, we measured approximately 2% of alleles with indels, demonstrating specificity of this nuclease for the N98S allele. In N98S iPSCs transfected with Sa.KKH RNP, we observed moderate rates of editing with approximately 10% of alleles having indels at the target site (Fig. 3B). In WT iPSCs transfected with Sa.KKH RNP, we observed no detectable indel generation, demonstrating that this nuclease is also specific for the N98S allele. Among the alleles edited by the Sp.HiFi RNP, we detected 2 categories of indel events: a 1 bp insertion and a 9 bp deletion (Fig. 3C). Among the alleles edited by the Sa.KKH RNP, we also detected 2 categories of indel events, but in this case a 9 bp deletion was predominant, followed by a 15 bp deletion (Fig. 3C).

**Figure 3:**
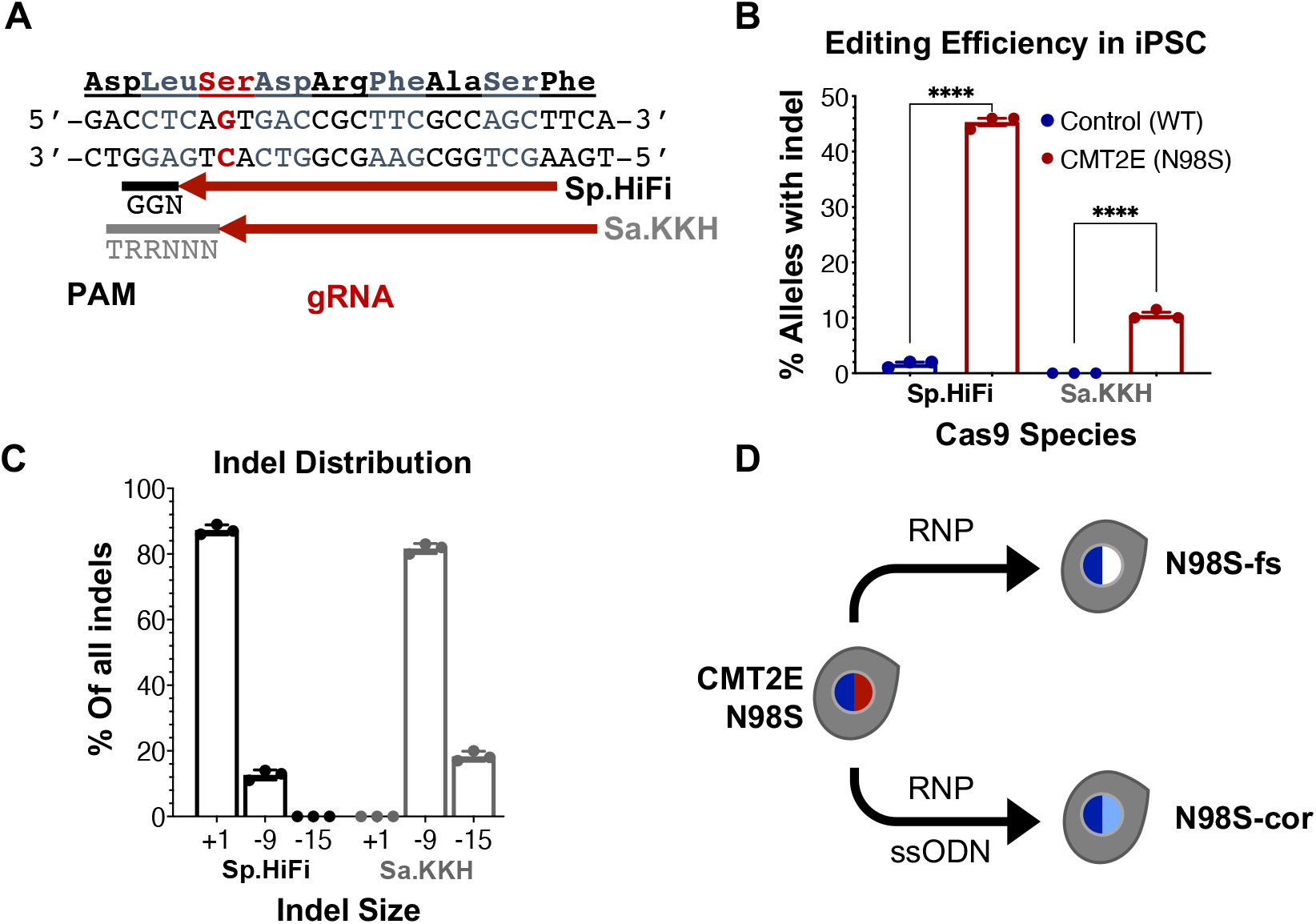
Allele-specific editing of *NEFL* N98S mutation in iPSCs. (A) *NEFL* target sequence with N98S mutation in red and gRNAs for Sp.HiFi and Sa.KKH Cas9 targeting the antisense strand. The canonical PAM recognition sequences for Sp.HiFi and Sa.KKH are shown in black and gray, respectively. (B) Efficiency and specificity of indel generation in control and CMT2E patient lines after transfection with Cas9/gRNA ribonucleoprotein (RNP), measured by ICE analysis, **** p < 0.0001 by t-test. (C) Comparison of indel sizes generated in CMT2E patient line by Cas9 enzyme. (D) Schematic of the derivation of clonal edited iPSC lines from the CMT2E N98S patient line. Transfection of N98S iPSC with Sp.HiFi RNP produced N98S-fs (frameshift inactivating the mutant allele). Transfection of N98S iPSC with Sp.HiFi RNP plus single-strand oligonucleotide donor (ssODN) produced N98S-cor (precise correction of the mutation). Colored nuclei indicate *NEFL* genotype: dark blue = wild type, red = N98S mutant, white = frameshift (knockout), light blue = correction of N98S to wild type.

The 9 and 15 bp deletion events are unlikely to disrupt expression of the N98S allele as they do not create a shift in reading frame. On the other hand, the 1 bp insertion, which constituted approximately 87% of the total indel events with Sp.HiFi N98S RNP, represents a potential therapeutic edit, as it results in a frameshift mutation. We therefore isolated a clone with a 1 bp insertion at the target site, referred to here as N98S-frameshift (fs). The frameshift mutation leads to a premature stop codon immediately after residue 98 of the N98S allele, which we predicted would effectively interrupt expression from this allele (Supplementary Fig. S3B). For comparison, we designed a strategy for precise correction of the N98S allele to a wild-type sequence by homology directed repair (HDR). To achieve this, we transfected N98S iPSCs with Sp.HiFi RNP and a single-strand DNA oligonucleotide donor (ssODN). The ssODN was designed to precisely repair the N98S mutation and insert a silent mutation to facilitate genotyping. The efficiency of this editing outcome was low (2.5%), so we utilized sib-selection to enrich for cells with the desired HDR event (Miyaoka et al., 2014) (Supplementary Fig. S3C). We isolated a clone, referred to as N98S-corrected (N98S-cor), with precise repair of the N98S sequence as well as insertion of silent mutation c.297C>T (Supplementary Fig. S3D).

### Frameshift indel reduces mutant *NEFL* mRNA and protein expression in N98S motor neurons

To investigate how allele-specific editing of the N98S allele affects *NEFL* mRNA and protein expression, we differentiated WT, N98S, N98S-fs, and N98S-cor iPSCs into i^3^LMNs. In addition, we generated i^3^LMNs from previously characterized iPSC lines with heterozygous (*NEFL* +/−) and homozygous (*NEFL* −/−) excision of the *NEFL* coding exons (Watry et al., 2020). *NEFL* +/− and *NEFL* −/− i^3^LMNs are isogenic to our WT control and were used as a comparison for *NEFL* mRNA and NF-L protein expression. We measured total *NEFL* mRNA levels in i^3^LMNs by quantitative RT-ddPCR and found that N98S-fs i^3^LMNs had 52% less *NEFL* mRNA than their unedited counterparts. By comparison, *NEFL* +/− i^3^LMNs had 36% less *NEFL* mRNA than WT i^3^LMNs (Fig. 4A). Thus, the frameshift introduced by the 1bp insertion disrupted expression of the edited allele as effectively as complete excision of the coding sequence. WT, N98S, and N98S-cor i^3^LMNs all exhibited similar levels of *NEFL* mRNA expression. To compare changes in transcription with protein levels, we measured NF-L protein by western blot in i^3^LMNs from the same panel of iPSC lines. Like *NEFL* mRNA levels, NF-L protein levels were reduced (by 42%) in N98S-fs i^3^LMNs compared to N98S i^3^LMNs (Fig. 4B). Interestingly, *NEFL* +/− i^3^LMNs had a non-significant reduction in NF-L protein compared to WT i^3^LMNs. The reason for the differing degrees of NF-L protein reduction in *NEFL* +/− and N98S-fs i^3^LMNs is not clear, but may be due to differences in genetic background and genetic compensation mechanisms. WT, N98S, and N98S-cor i^3^LMNs exhibited similar levels of total NF-L protein. Thus, the accumulation of NF-L protein in the cell bodies of N98S i^3^LMN is consistent with a defect in localization and intracellular transport, rather than perturbations in protein synthesis or degradation. Neither *NEFL* mRNA nor NF-L protein was detected in the *NEFL* −/− samples, validating the specificity of our assays.

**Figure 4:**
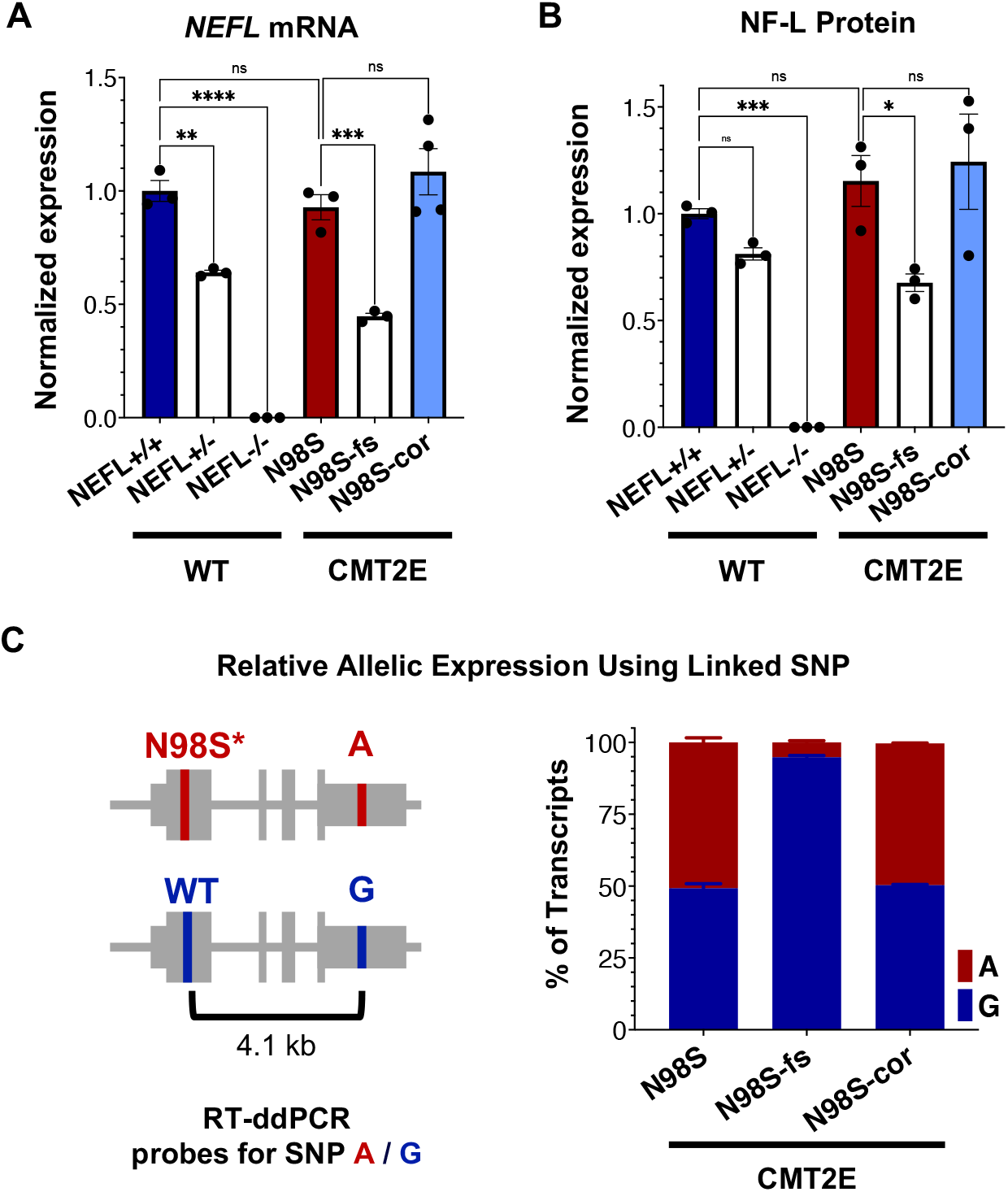
*NEFL* expression in allelic series of edited motor neurons. A series of edited iPSCs derived from a healthy control (WT) and CMT2E patient were differentiated into i^3^LMNs, and RNA was extracted on day 7 or protein on day 10. (A) Total *NEFL* transcript expression by quantitative RT-ddPCR, normalized to *GAPDH*. (B) Total NF-L protein expression by western blot, normalized to GAPDH. (C) Relative expression of *NEFL* alleles in i^3^LMNs derived from CMT2E patient line and edited subclones. Allele-specific RT-ddPCR targeting a heterozygous SNP in the 3’ UTR was used to genotype transcripts using linkage shown in the schematic. All bar graphs represent mean +/− S.E.M. For all graphs ns = not significant, * p < 0.05, ** p < 0.01, *** p < 0.001, **** p < 0.0001 by one-way ANOVA with Šídák’s test for multiple comparisons.

To confirm that the observed decrease in total *NEFL* expression in N98S-fs i^3^LMNs was specific to the mutant allele, we measured the relative expression of the mutant and wild-type alleles in N98S, N98S-fs, and N98S-cor i^3^LMNs. Allele specificity was achieved by targeting a RT-ddPCR assay to a heterozygous SNP present in the 3’ untranslated region of the *NEFL* gene in our CMT2E patient line (Fig. 4C). In N98S and N98S-cor i^3^LMNs, both alleles were transcribed at equal rates, with each representing 50% of the total transcripts detected. However, in N98S-fs i^3^LMNs, *NEFL* mutant allele transcripts constituted only 5% of the total *NEFL* transcripts detected, suggesting that the premature stop codon results in efficient nonsense mediated decay of the mutant mRNA. Overall, these experiments demonstrate that the frameshift mutation leads to dramatic reduction of mutant *NEFL* mRNA and, presumably, mutant NF-L protein in i^3^LMNs, while isogenic correction of the N98S mutation with introduction of a silent mutation does not alter *NEFL* expression.

### Selective inactivation of the N98S allele rescues pathologic phenotypes in motor neurons

To determine if inactivation of the *NEFL* mutant allele prevents the pathologic phenotype, we measured the distribution of NF-L in fixed and stained WT, N98S, N98S-fs, and N98S-cor i^3^LMNs (Fig. 5A-C). To increase the throughput of this analysis, we developed an automated image analysis pipeline using CellProfiler to identify i^3^LMN cell bodies and quantify NF-L intensity (Fig. 5C, Supplementary Fig. S5). Consistent with the manually processed data, N98S i^3^LMN cell bodies displayed significantly higher NF-L fluorescence intensity values compared to WT. N98S-fs i^3^LMNs displayed significantly lower NF-L fluorescence intensity values than N98S i^3^LMNs, demonstrating that inactivation of the mutant *NEFL* allele prevents the pathologic accumulation of NF-L in the cell bodies of motor neurons. In addition, the NF-L fluorescence intensity values of N98S-fs i^3^LMNs were indistinguishable from those of WT and N98S-cor i^3^LMNs, further suggesting that NF-L localization and transport is normalized after inactivation of the N98S mutant allele. We also investigated whether gene editing altered the ability of the neurons to generate functional action potentials. MEA analysis demonstrated that the N98S-fs and N98S-cor i^3^LMNs retained similar spontaneous electrical activity as their unedited N98S counterparts (Supplementary Fig. S7).

**Figure 5:**
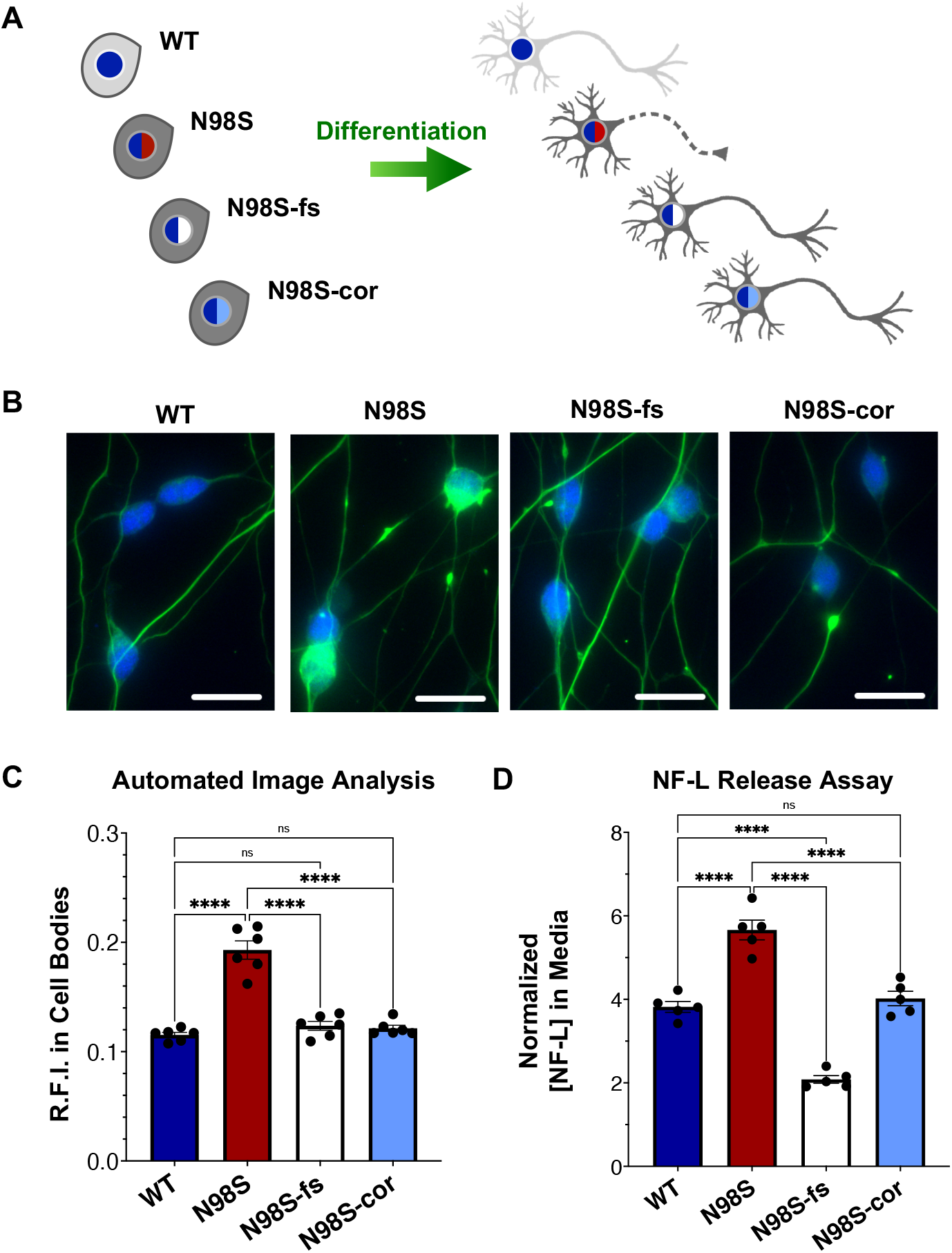
Rescue of pathologic phenotypes in edited N98S motor neurons. (A) Schematic of allelic series of iPSCs differentiated into healthy (continuous axon) or diseased (dashed axon) i^3^LMNs. Colored nuclei represent *NEFL* genotypes as described in figure 3. Cytoplasmic shading indicates differential genetic backgrounds, dark grey cells are isogenic. (B) Representative images of day-7 i^3^LMNs stained with anti-NF-L (green) and anti-HB9 (blue) antibodies. Scale bars = 25 μM. (C) Quantification of NF-L relative fluorescence intensity (R.F.I.) in HB9+ cell bodies using automated image analysis pipeline. Data points represent mean values from independent differentiations, bars indicate mean of six biological replicates +/− S.E.M. (D) ELISA for NF-L protein in media from day-14 i^3^LMNs. NF-L levels were normalized to neurite density measurements to correct for well-to-well variability in cell seeding (Supplementary Fig. S6). Individual data points shown, bars indicate mean of five biological replicates +/− S.E.M. For all graphs **** p < 0.0001 by one-way ANOVA with Šídák’s test for multiple comparisons.

Finally, we assayed NF-L release in the culture media as a means to assess cellular integrity of our i^3^LMNs. NF-L is a biomarker of neurodegenerative diseases, as NF-L levels in cerebrospinal fluid and serum reflect the extent of neuroaxonal damage leading to neurofilament release (Lu et al., 2015; Hansson et al., 2017; Disanto et al., 2017). Correspondingly, it has been reported that iPSC-derived motor neuron cultures with the *NEFL*-N98S mutation release more NF-L into the media than healthy controls, consistent with increased axonal degeneration (Maciel et al., 2020). Using a commercially available enzyme-linked immunosorbent assay (ELISA), we measured the levels of NF-L in media from day-14 i^3^LMN cultures (Fig. 5D). As expected, media from N98S i^3^LMNs had significantly higher normalized levels of NF-L than media from WT i^3^LMNs, suggesting that the N98S mutation results in neuroaxonal damage. N98S-cor and WT i^3^LMN cultures exhibited equivalent levels of NF-L, confirming that the increase in NF-L levels observed in N98S i^3^LMNs was due to the N98S mutation rather than inherent cell line variability. In contrast, N98S-fs i^3^LMNs released significantly less NF-L than all other cell lines, including the WT and N98S-cor controls. This was expected, as overall NF-L expression is reduced in N98S-fs i^3^LMNs (Fig. 4B). To determine whether this observation was simply due to changes in intracellular protein expression, we compared cell lines by differences in NF-L released into media (as measured by ELISA, Fig. 5D) versus differences in intracellular NF-L levels (as measured by western blot, Fig. 4B). The relative ratios of N98S-fs to N98S-cor were similar; 0.52 for NF-L released in media and 0.55 for intracellular NF-L. This demonstrates that the differences in released NF-L can be completely explained by intracellular protein levels and suggests equivalent degrees of neuronal integrity when comparing the two editing strategies. Conversely, the relative ratios of N98S-fs to parental N98S were 0.37 for NF-L released in media and 0.59 for intracellular NF-L. The larger decrease in NF-L released into the media supports the conclusion that inactivation of the mutant allele leads to both decreased NF-L protein expression, as well as improved neuronal integrity compared to the unedited diseased neurons.

In summary, we conclude that inactivation of the mutant allele by CRISPR-Cas9 gene editing effectively prevents disease phenotypes in i^3^LMNs derived from a patient with autosomal dominant CMT2E caused by the N98S mutation. Moreover, N98S-fs i^3^LMNs were phenotypically similar to N98S-cor i^3^LMNs, suggesting that knockout of the mutant allele is as effective as precise repair of the mutation in this model system.

## DISCUSSION

In this study, we used allele-specific gene editing with engineered Cas9 enzymes to inactivate a missense mutation causing CMT2E, a severe autosomal dominant neuromuscular disease. Cas9-induced inactivation was efficient, selective, and effectively reversed the accompanying pathology in patient iPSC-derived motor neurons without apparent adverse effects. In the process, we incorporated rapid neuronal differentiation and high-throughput automated image analysis to enable measurement of changes in disease phenotypes in a fraction of the time required by previous methodology. Our findings establish the framework for therapeutic development of gene editing for CMT2E and for other motor neuron diseases caused by dominant missense mutations.

### Gene editing as an efficient path to allele-specific gene inactivation

We selected CRISPR-Cas9-based genome editing because it presents several advantages over other therapeutic strategies, with important clinical implications (Porteus, 2019). First, the therapeutic construct only needs to be active briefly and requires only a single treatment, since the outcome is a permanent change in the cellular genome. In fact, transient expression is desirable as it minimizes the risk of off-target effects and immune response to the vector construct. Second, since it targets the mutation itself, gene editing can achieve complete correction or inactivation of the mutant allele in treated cells, rather than mere attenuation of the mutation’s effect. Finally, even single-nucleotide mutations can be targeted by Cas nucleases with a high degree of specificity (Wienert et al., 2019), suggesting the approach could be safely deployed to treat a large number of conditions caused by missense mutations.

We demonstrated that Cas9 enzymes derived from both *Staphylococcus aureus* and *Streptococcus pyogenes* can induce indels specifically at the *NEFL*-N98S mutant allele, and that the latter predominantly leads to a 1-bp insertion that efficiently disrupts mutant allele expression. In contrast, the former preferentially generates 9- and 15-bp deletions that are less useful for disrupting expression of the mutant allele. Interestingly, 9- and 15-bp deletions are predicted by microhomology domains near the cleavage sites for both of our Cas9 nucleases (Bae et al., 2014). Large-scale studies have begun to define the mutational profiles produced by Cas9-induced cleavage and predict outcomes produced by specific gRNA, but our findings highlight how these predictions are not necessarily transferable to alternate enzymes, which differ in the nature of the double-strand breaks produced (Chen et al., 2019; Allen et al., 2018; Tycko et al., 2018). Finally, these predictions are based on experiments performed in immortalized cell lines and the generalizability to primary and post-mitotic cells is yet to be determined.

We achieved phenotypic rescue using CRISPR gene editing both by generating indels in the *NEFL* allele carrying the N98S mutation and by correcting the N98S mutation to wild type via HDR. In both cases, gene editing starts with the creation of targeted double-strand breaks, but the outcome depends on the endogenous DNA repair machinery that repairs the break—non-homologous end-joining (NHEJ) or homology-directed repair (HDR) (Porteus, 2019). NHEJ is most useful for gene inactivation via frameshift-inducing indels, as in our approach, and is active even in post-mitotic neurons (Bétermier et al., 2014; Shalem et al., 2014; Gaj et al., 2017). Precise correction of a mutant sequence by HDR requires a donor template to facilitate homologous recombination but is typically much less efficient than NHEJ and active only in proliferative cells (Miyaoka et al., 2016). Indeed, in our experiments editing outcomes from NHEJ were over 10 times more efficient than HDR using the same gRNA and Cas9 enzyme. Both approaches were able to effectively rescue disease phenotypes in CMT2E i^3^LMNs, but as HDR is not active in post-mitotic cells like neurons, inactivation of the mutant allele via NHEJ is the more tractable therapeutic strategy. An important caveat is that allele-specific gene inactivation is restricted to genes in which heterozygous loss of function is well tolerated, as is the case for *NEFL*.

### Alternative strategies and future directions

We validated CRISPR reagents that are specific to the human *NEFL*-N98S mutation, but there are a multitude of distinct missense mutations in *NEFL* that cause CMT2E (Stone et al., 2019). The majority occur in the first exon and should be amenable to the same editing strategy that we employed here. However, inducing frameshifts at the site of mutations in distal coding exons may not be therapeutic, as nonsense codons in that context can lead to expression of a pathologic truncated protein rather than nonsense-mediated decay (Lindeboom et al., 2016; Stone et al., 2019). Alternate editing strategies for these mutations include excision of mutant alleles using pairs of guide RNAs, or precise correction using non-nuclease editing technologies such as base editors or PRIME editing (Watry et al., 2020; Komor et al., 2016; Anzalone et al., 2019).

Alternative gene therapy approaches include gene augmentation, more commonly used to treat recessive loss-of-function mutations, and RNA interference; however, both present their own challenges. Gene augmentation in dominant disease can have partial benefits (Mao et al., 2011) yet it requires high levels of exogenous gene expression to overcome effects of the mutant allele, and will be ineffective if the mutant gene product continues to interfere with the function of the wild-type gene product, or causes direct toxicity to the cell (Yu-Wai-Man, 2016; Diakatou et al., 2019). RNA interference, where a vector delivers a small RNA that targets the mutant transcript for degradation, has been used successfully in animal models of dominant disease, including CMT2D (Matsa et al., 2014; Zaleta-Rivera et al., 2019; Morelli et al., 2019). Although promising, this approach needs to overcome significant challenges including efficiency and specificity, particularly for single-nucleotide point mutations. Furthermore, both gene augmentation and RNA interference require long-term persistence of the vector or repeated treatments.

The major limitation of our studies is that we performed editing in stem cells rather than in the neurons directly. Efficient delivery of gene-editing reagents to neurons in vitro is challenging, and ongoing work in our group and others is focused on improving methodology to enable routine gene-editing experiments in iPSC-derived neurons. The studies presented here establish a crucial baseline, by identifying candidate therapeutic reagents and validating rapid development of disease-specific phenotypes that can be assayed in high throughput. These tools will be invaluable to answer outstanding questions including the applicability to other missense mutations in *NEFL*, allele specificity and repair outcomes in neurons, and the time course for reversing the established disease phenotype. Furthermore, in vivo animal experiments will be important to complement the in vitro human models. Fortunately, in the case of CMT2E there is an excellent animal model in the N98S knock-in mouse that precisely mimics the human mutation, and is associated with a severe and well characterized phenotype (Adebola et al., 2015; Lancaster et al., 2018). In the case of gene editing, the ultimate therapeutic reagent must be primarily developed in a human system due to species-specific genome sequences, but in vivo experiments will be critical to optimize delivery methods, assess functional outcomes, and evaluate systemic side effects.

## Author Contributions

LJ, MS, and BC conceived the project. CF, KW, HW, and LJ designed the experiments. CF performed the majority of experiments and data analysis with assistance from other authors as specified. KW, HG, AL, and HW transfected iPSC and isolated clonal lines. KW optimized neuron differentiation, western blots, and immunofluorescent staining. CM performed selected gene editing and immunofluorescent staining. CM, LZ, and TM conducted multielectrode array analysis. GR helped optimize neuron differentiation and design of the CellProfiler analysis pipeline. CF and LJ wrote the manuscript with assistance from all authors.

## Acknowledgements

We thank Michael Ward for providing the AAVS1-hNIL iPSC line and AAVS1-hNIL and CLYBL-hNIL plasmids, the Gladstone Stem Cell Core and Cell Line Genetics for their services, and A. Birk, A. Sachdev, Z. Nevin, C. Clelland, and F. Chanut for technical assistance, advice, and critical review of the manuscript.

