## Supplementary Figures and Tables for "Allele-specific gene editing rescues pathology in a human model of Charcot-Marie-Tooth disease type 2E"

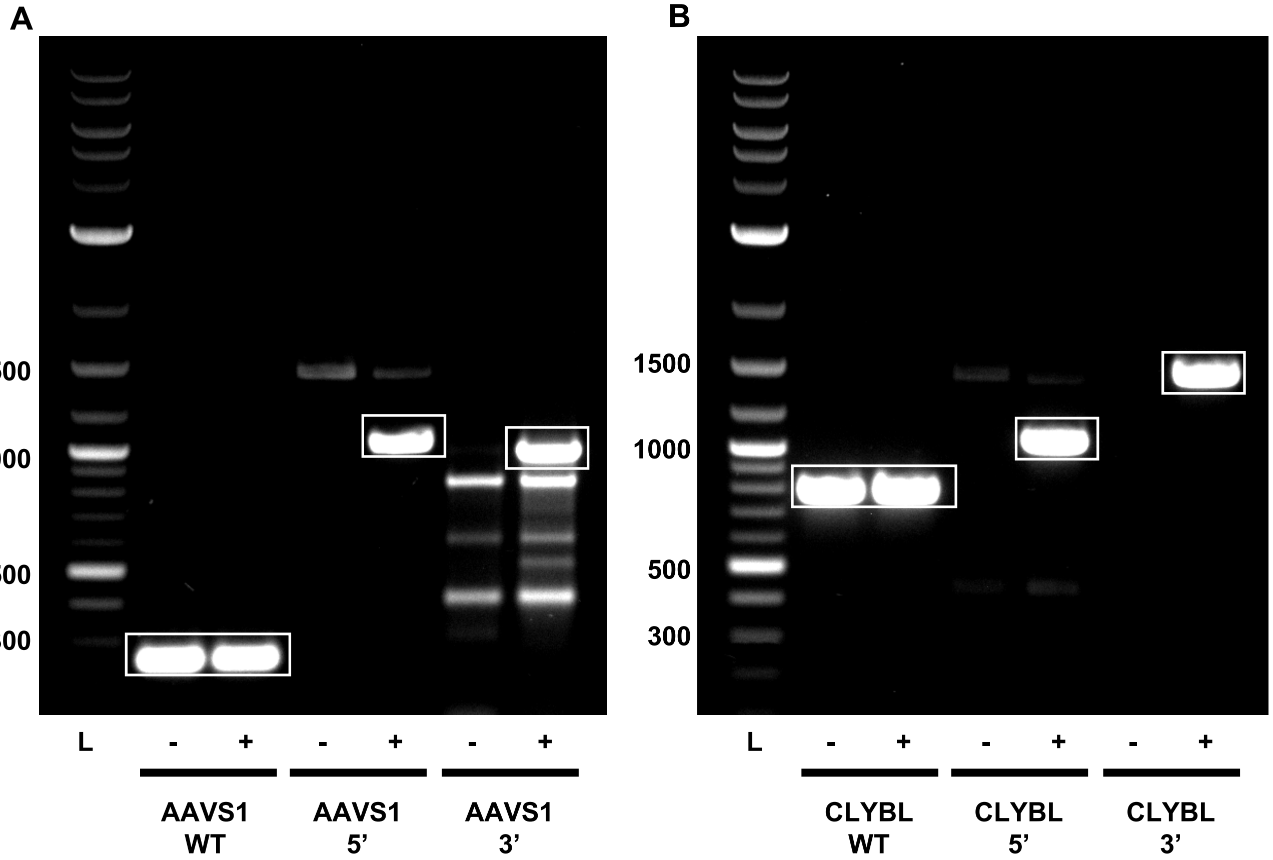


**Supplementary Figure S1: Confirmation of targeted integration of hNIL cassette in iPSC lines.** PCR assays were designed to amplify the unmodified AAVS1 or CLYBL locus (WT) as well as the left (5’) and right (3’) integration junctions. Genomic DNA from unmodified iPSC (-) was compared to genomic DNA from clonal lines isolated after transfection with TALENs and hNIL vector (+). L = ladder. (A) AAVS1 integration of hNIL in CMT2E-N98S line. (B) CLYBL integration of hNIL in the WT line. Boxes indicate amplicon products matching the expected size for each assay as follows:

WT AAVS1 = 254 bp
AAVS1 hNIL 5' junction = 992 bp
AAVS1 hNIL 3' junction = 989 bp
WT CLYBL = 790 bp
CLYBL hNIL 5' junction = 1032 bp
CLYBL hNIL 3' junction = 1470 bp

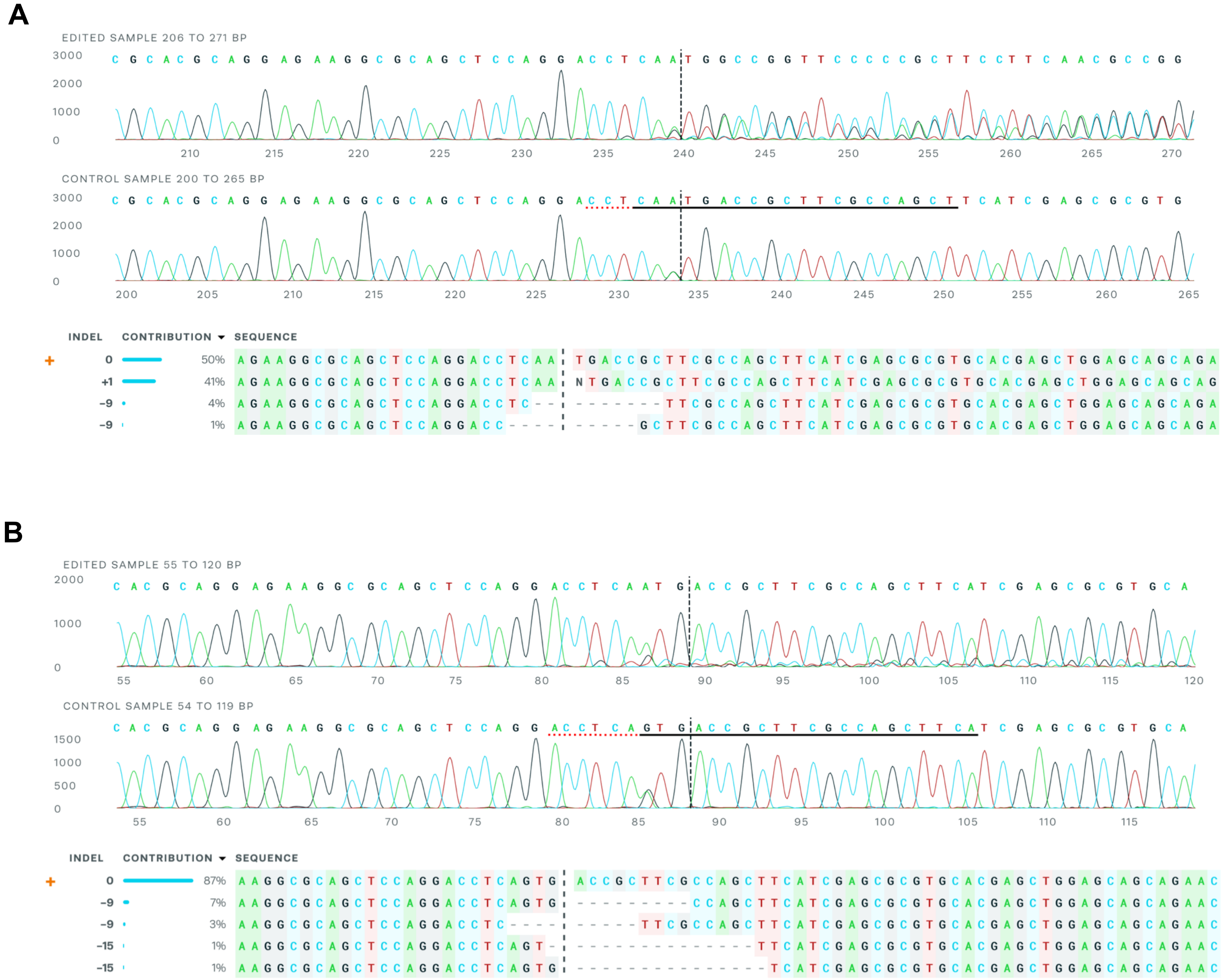


**Supplementary Figure S2: Representative ICE analyses for editing with Sp. and Sa.KKH Cas9.** Sequencing chromatograms from samples edited with Sp.HiFi (A) and Sa.KKH Cas9 (B) are shown along with unedited controls. The gRNA target sequences are indicated with black solid underlines, PAM sequences with red dotted underlines, and predicted cut sites with vertical dashed lines. The identities of various detected indel outcomes are shown below, with contribution as a percentage of all alleles. Unedited alleles are marked with an orange +.

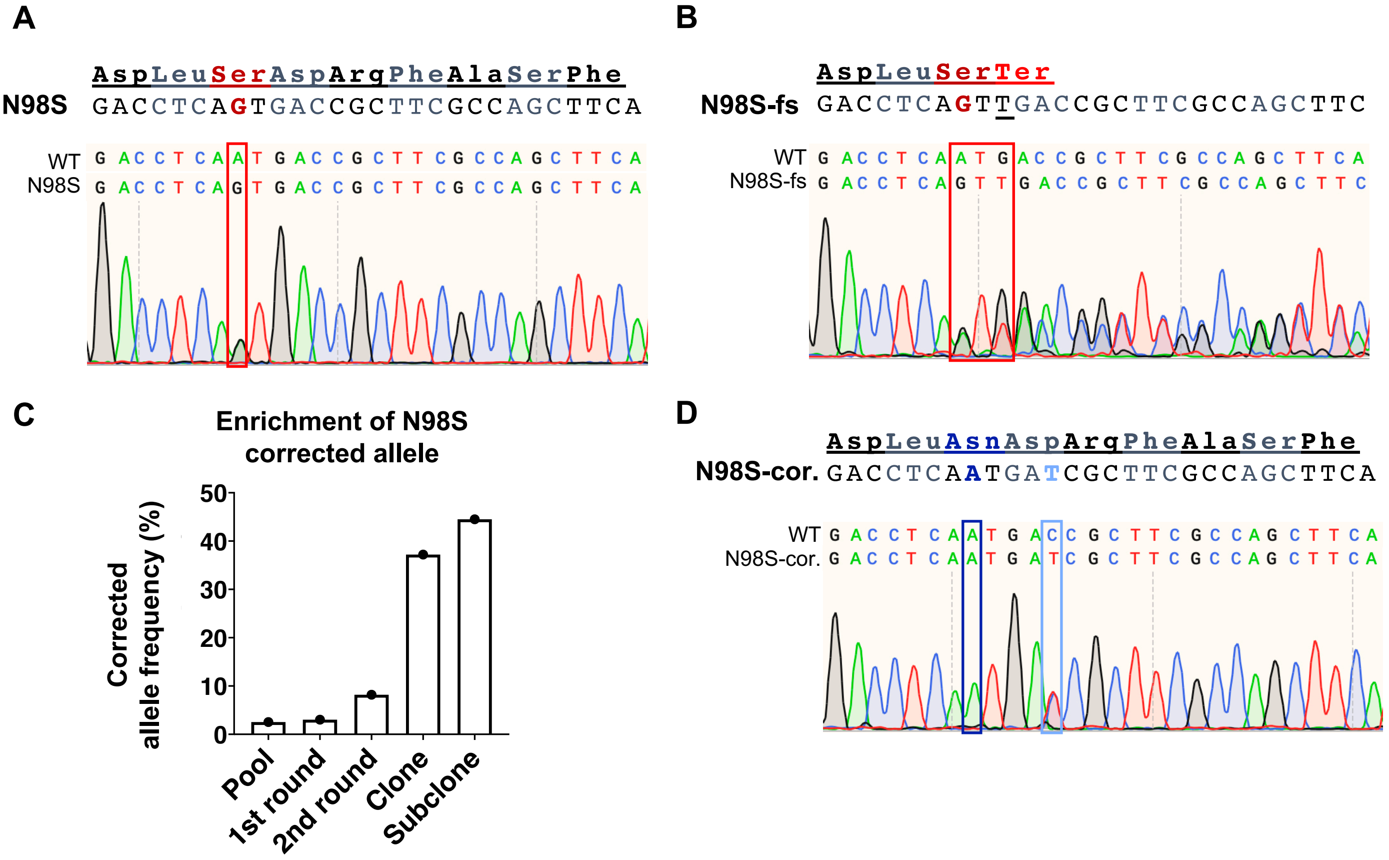


**Supplementary Figure S3: Isolation and genotyping of edited CMT2E-N98S iPSC lines.** (A) Sequence of N98S mutant allele. Sequencing chromatogram below from *NEFL*-N98S iPSC line with the heterozygous N98S mutation boxed in red. (B) Sequence of N98S-fs allele demonstrating insertion of one extra thymidine nucleotide (T) leading to immediate nonsense codon (Ter). Sequencing chromatogram from the N98S-fs clone below, with the site of mutation and +1bp insertion boxed in red. (C) Serial sib selection led to enrichment of the N98S corrected allele, followed by manual clone picking. (D) Sequence of N98S-cor. allele with G->A substitution to correct disease mutation along with C->T silent mutation to facilitate genotyping. Sequencing chromatogram from the N98S-cor clone below, with the corrected mutation site boxed in dark blue and the silent SNP boxed in light blue.

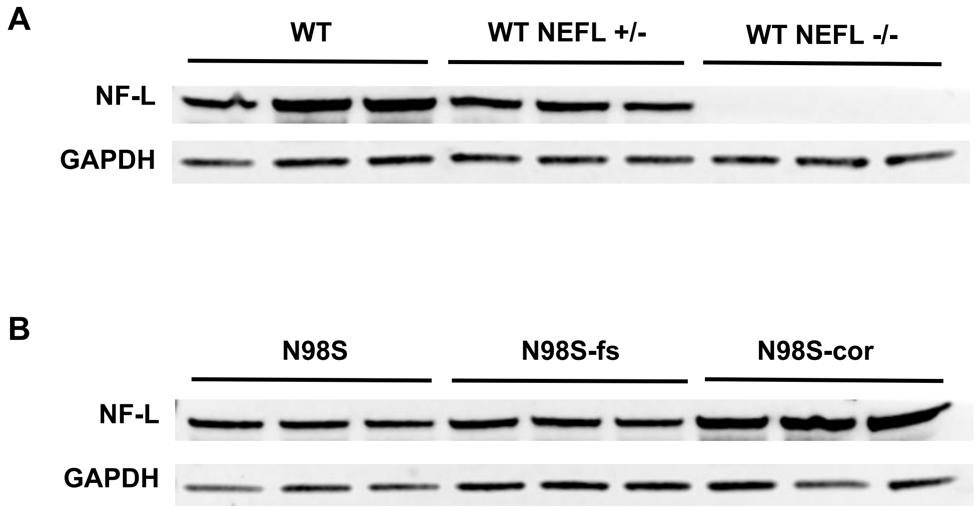


**Supplementary Figure S4: Western blots for quantification of NF-L protein levels.** Total protein was extracted from day 10 i$^3$LMNs, blots probed with anti-neurofilament light (NF-L) and anti-GAPDH antibodies**.** Quantification is shown in figure 4. (A) Samples derived from healthy control (WT) neurons and isogenic lines with heterozygous or homozygous deletion of *NEFL*. (B) Samples derived from CMT2E N98S neurons and isogenic lines edited with frameshift mutation of the N98S allele (N98S-fs) or correction of N98S to wild type sequence (N98S-cor).

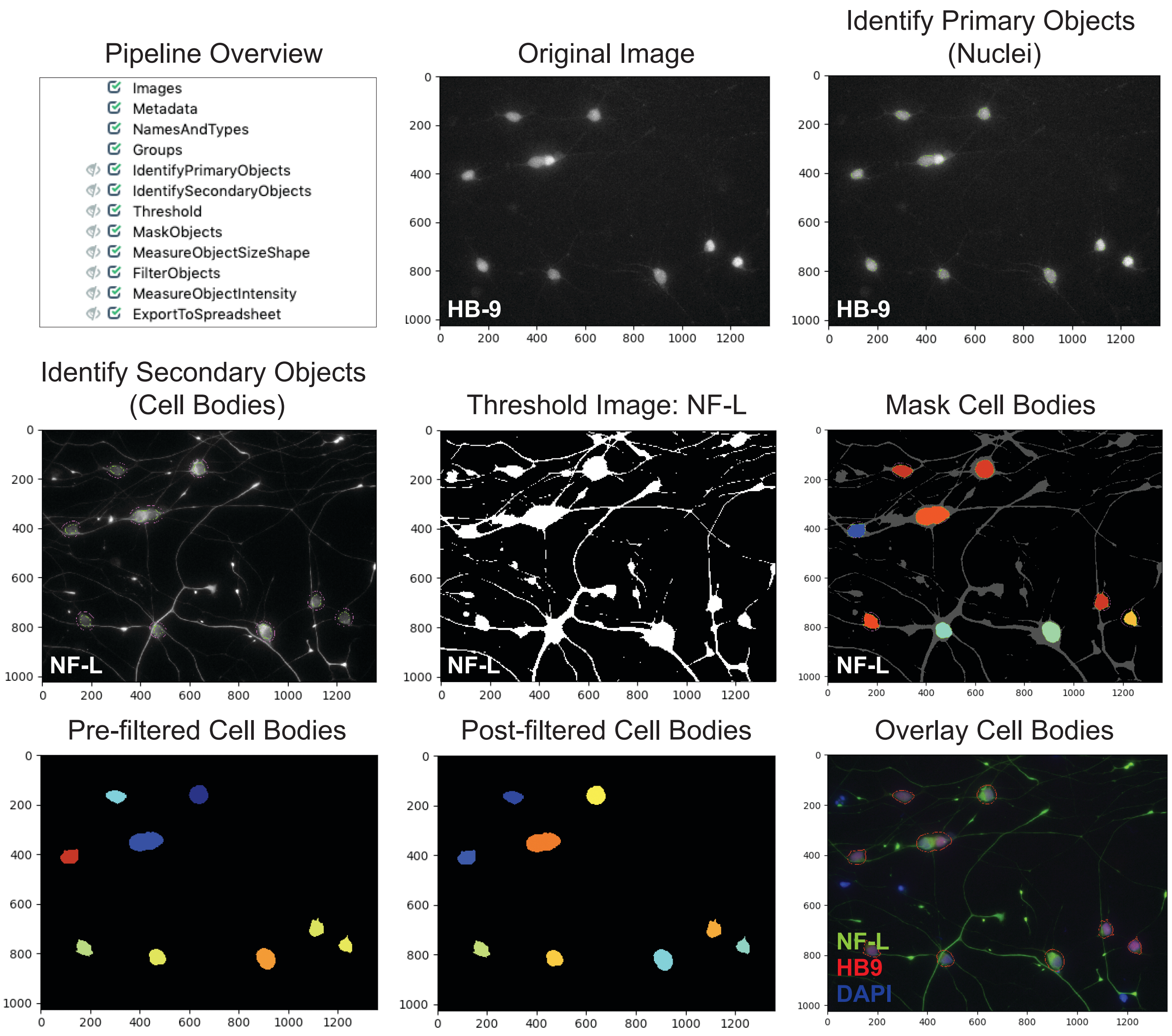


**Supplementary Figure S5: CellProfiler pipeline for quantification of NF-L fluorescence intensity in motor neuron cell bodies.** HB9+ nuclei (primary objects) were identified. Cell bodies (secondary objects) were identified by extending the area of HB9+ radially by 15 pixels. The image with NF-L staining was thresholded and used to mask the cell bodies to exclude areas containing only background. The cell bodies were then filtered by shape and compactness to remove any incorrectly identified objects. The average NF-L intensity of the cell bodies was measured.

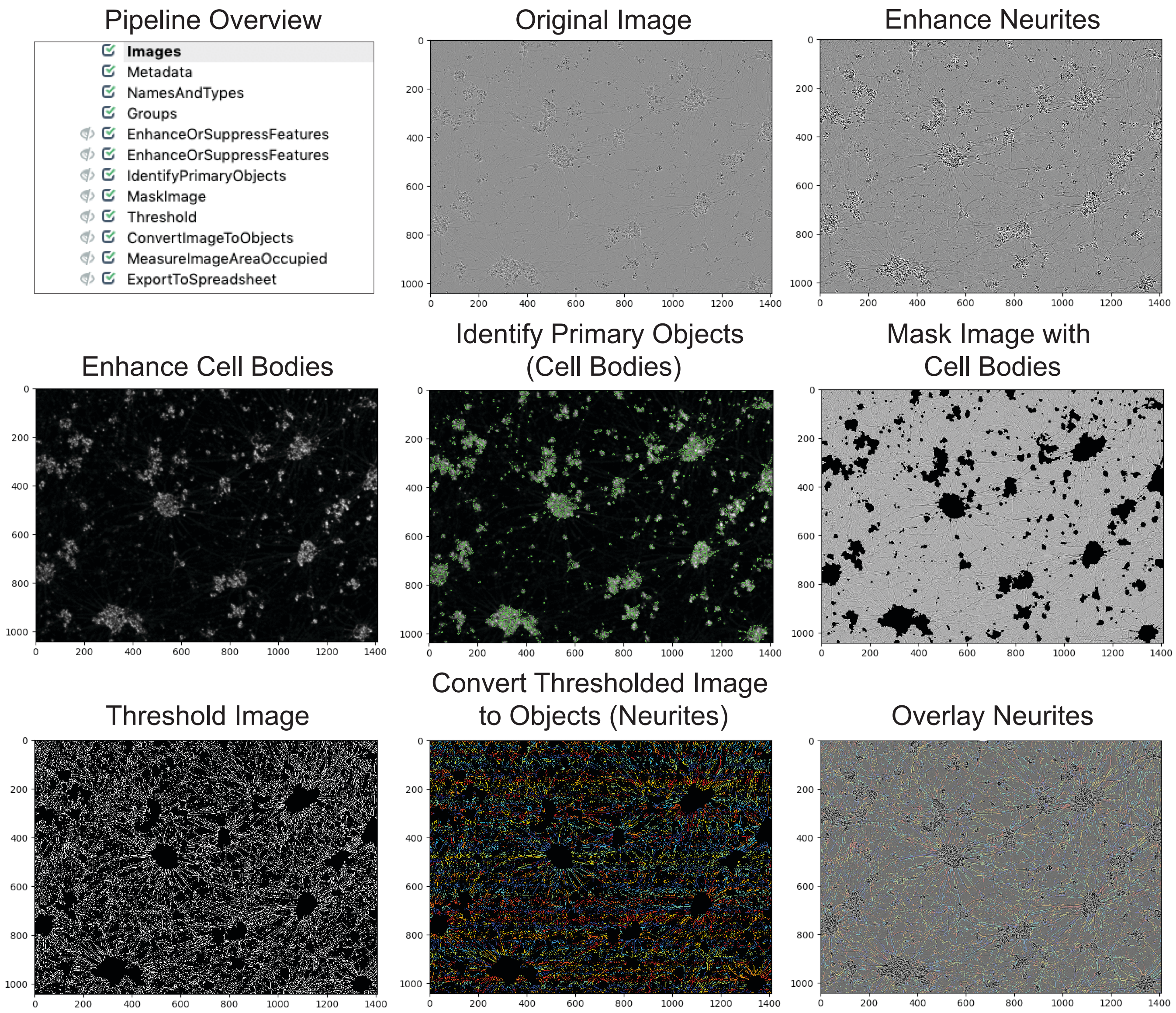


**Supplementary Figure S6: CellProfiler pipeline for quantification of neurite density.** Neurites and cell bodies were enhanced by the EnhanceOrSuppressFeatures module. Cell bodies were identified and removed from the image. The image was thresholded to distinguish the neurites from the background. The thresholded image was then used to identify the neurites as objects. The total area occupied by the neurites was measured. NF-L protein levels in media assessed by ELISA assay were normalized to the measurement of neurite density to correct for well-to-well variability in cell density.

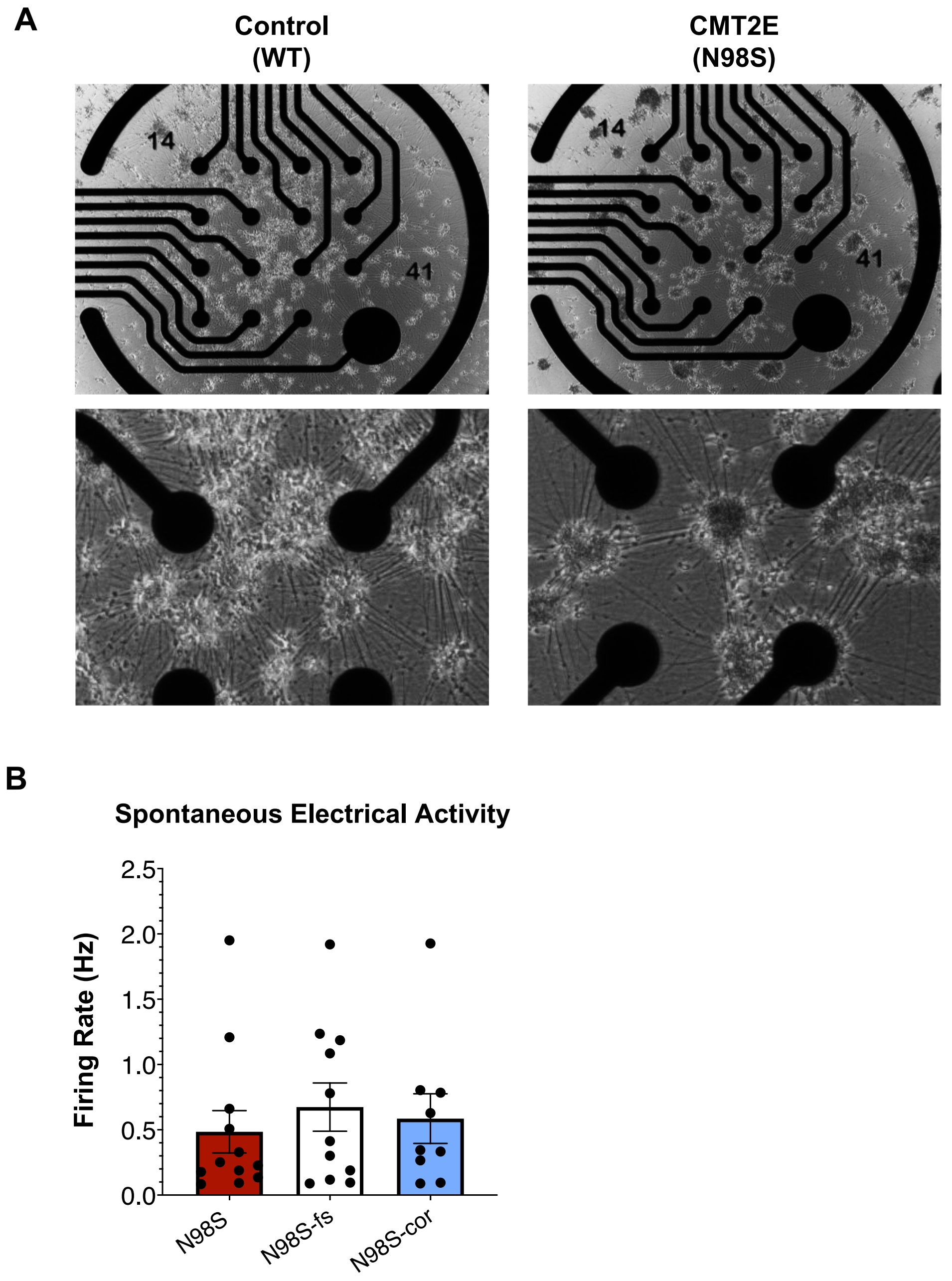


**Supplementary Figure S7: MEA analysis in motor neurons.** (A) Representative images of control and CMT2E neurons on MEA plates. Images were acquired on differentiation day 14. (B) Measurements of spontaneous electrical activity in edited $i^3$LMN measured by mean firing rate of action potentials detected by MEA. Cells were seeded on MEA plates on differentiation day 3 and recordings were obtained on differentiation day 14. Each data point represents the weighted firing rate from an individual well, bars mean of all wells +/- S.E.M.

**Supplementary Table 1: Oligonucleotide Sequences**

| **Name** | **Sequence (5' → 3')** |
|---|---|
| hNIL AAVS1 5' Junction F | CCTGAGTCCGGACCACTTTG |
| hNIL AAVS1 5' Junction R | AGAAGACTTCCTCTGCCCTC |
| hNIL AAVS1 3' Junction F | GCCTGGTAGACAGGGCTGG |
| hNIL AAVS1 3' Junction R | TGTGGGGTGGAGATATCAGC |
| AAVS1 WT F | CGGTTAATGTGGCTCTGGTT |
| AAVS1 WT R | AGGATCCTCTCTGGCTCCAT |
| hNIL CLYBL 5' Junction F | CAGACAAGTCAGTAGGGCCA |
| hNIL CLYBL 5' Junction R | AGAAGACTTCCTCTGCCCTC |
| hNIL CLYBL 3' Junction F | CACCAGCAACCTGACGTTTT |
| hNIL CLYBL 3' Junction R | TTTTATAGGCGCCCACCGTA |
| CLYBL WT F | TGACTAAACACTGTGCCCCA |
| CLYBL WT R | AGGCAGGATGAATTGGTGGA |
| N98S Sequencing F | TACTCGACCTCCTACAAGCG |
| N98S Sequencing R | GCGGCTCTTGAACCATTCCT |
| ssODN for HDR | CACGCGCTCGATGAAGCTGGCGAAGCG<u>A</u>TCA<u>T</u>TGAGGTCCTGGAGCTGCGCCTTCTCCTG |

**Supplementary Table 2: Guide RNA Sequences**

| Name | gRNA spacer sequence |
|---|---|
| N98S Sp.HiFi | AGCTGGCGAAGCGGTCACTG |
| N98S Sa.KKH | TGAAGCTGGCGAAGCGGTCAT |

**Supplementary Table 3: Antibodies**

| Name/Antigen | Host | Vendor/Catalog # | IF Concentration | WB Concentration |
|---|---|---|---|---|
| β-Tubulin III | Rabbit | Sigma T2200 | 1:500 | |
| SMI33 (NF-H/M) | Mouse | Biolegend 835404 | 1:500 | |
| HB9 | Mouse | DSHB 81.5C10 | 1:200 | |
| NF-L | Rabbit | Sigma AB9568 | 1:500 | |
| NF-L | Mouse | Sigma N5139 | | 1:1000 |
| GAPDH | Rabbit | Abcam AB9485 | | 1:2000 |
| Anti-rabbit Alexa Fluor 488 | Goat | Invitrogen A11008 | 1:500 | |
| Anti-mouse Alexa Fluor 594 | Goat | Invitrogen A11005 | 1:500 | |
| Anti-rabbit IRDye 800CW | Donkey | Li-Cor 926-32213 | | 1:10,000 |
| Anti-mouse IRDye 680LT | Donkey | Li-Cor 926-68022 | | 1:10,000 |

**Supplementary Table 4: Gene Expression Assay IDs**

| Gene Target | Vendor | Primer/Probe Assay ID |
|---|---|---|
| *MNX1* | ThermoFisher | Hs00907365_m1 |
| *CHAT* | ThermoFisher | Hs00758143_m1 |
| *NEFL* | ThermoFisher | Hs01034882_m1 |
| *GAPDH* | Bio-Rad | qHsaCEP0041396 |
